# IPSC-derived midbrain astrocytes from Parkinson’s disease patients carrying pathogenic *SNCA* mutations exhibit alpha-synuclein aggregation, mitochondrial fragmentation and excess calcium release

**DOI:** 10.1101/2020.04.27.053470

**Authors:** Peter A. Barbuti, Paul Antony, Gabriella Novak, Simone B. Larsen, Clara Berenguer-Escuder, Bruno FR. Santos, Francois Massart, Dajana Grossmann, Takahiro Shiga, Kei-ichi Ishikawa, Wado Akamatsu, Steven Finkbeiner, Nobutaka Hattori, Rejko Krüger

**Affiliations:** Clinical and Experiment Neuroscience, Luxembourg Centre for Systems Biomedicine, University of Luxembourg, L-4362 Luxembourg; Transversal Translational Medicine, Luxembourg Institute of Health, L-1445, Luxembourg; Gladstone Institutes, the Taube/Koret Center for Neurodegenerative Disease, San Francisco, USA; Center for Genomics and Regenerative Medicine, Juntendo University School of Medicine, Japan; Department of Neurology, Juntendo University School of Medicine, Japan; University of California, San Francisco, USA; Parkinson Research Clinic, Centre Hospitalier de Luxembourg (CHL), Luxembourg

**Keywords:** midbrain astrocytes, patient-derived, Parkinson’s disease, *SNCA*, alpha-synuclein, mitochondria, calcium

## Abstract

Parkinson’s disease (PD) is characterized by the loss of A9 midbrain dopaminergic neurons and the accumulation of alpha-synuclein aggregates in remaining neurons. Many studies of the molecular and cellular basis of neurodegeneration in PD have made use of iPSC-derived neurons from patients with familial PD mutations. However, approximately half of the cells in the brain are glia, and their role facilitating neurodegeneration is unclear. We developed a novel serum-free protocol to generate midbrain astrocytes from patient-derived iPSCs harbouring the pathogenic p.A30P, p.A53T mutations in *SNCA*, as well as duplication and triplication of the *SNCA* locus. In our cellular model, aggregates of alpha-synuclein occurred only within the GFAP^+^ astrocytes carrying the pathogenic *SNCA* mutations. Assessment of spontaneous cytosolic calcium (Ca^2+^) release using Fluo4 revealed that *SNCA* mutant astrocytes released excess Ca^2+^ compared to controls. Unbiased evaluation of 3D mitochondrial morphometric parameters showed that these *SNCA* mutant astrocytes had increased mitochondrial fragmentation and decreased mitochondrial connectivity compared to controls, and reduced mitochondrial bioenergetic function. This comprehensive assessment of different pathogenic *SNCA* mutations derived from PD patients using the same cellular model enabled assessment of the mutation effect, showing that p.A53T and triplication astrocytes were the most severely affected. Together, our results indicate that astrocytes harbouring the familial PD mutations in *SNCA* are dysfunctional, suggesting a contributory role for dysfunctional astrocytes in the disease mechanism and pathogenesis of PD.

**Table of Contents Image:** 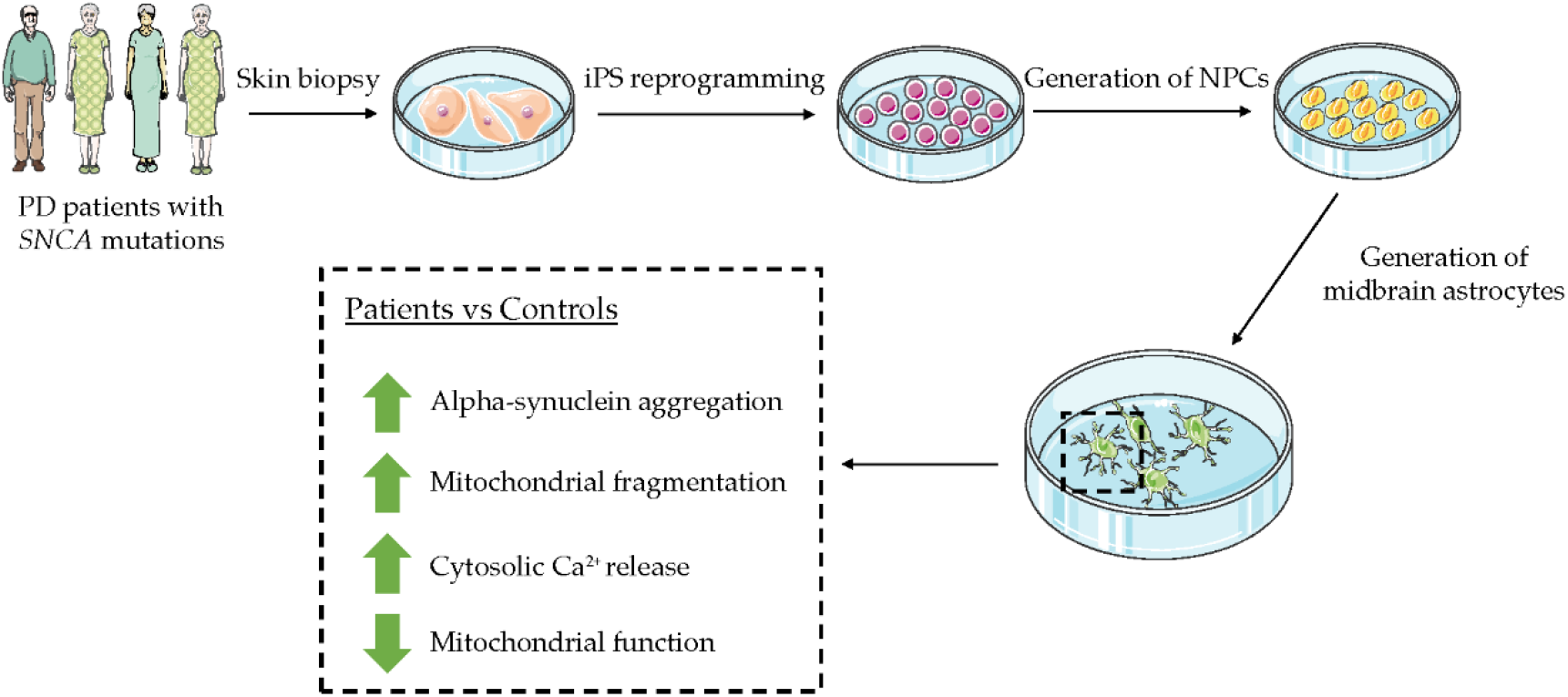

**Main Points:**

- We used a novel serum-free protocol to generate midbrain-specific functional astrocytes from Parkinson’s disease patients carrying pathological mutations in *SNCA*
- Patient-derived astrocytes show morphological and functional impairments

## Introduction

Parkinson’s disease (PD) is a neurodegenerative disease with two neuropathological hallmarks: the degeneration of neurons from the substantia nigra pars compacta (SNc) in the midbrain projecting to the striatum, and the accumulation of intracellular protein inclusions (Lewy bodies, LBs) in the neurons that remain. LBs are immunopositive for the alpha-synuclein protein, and contain crowded organelles and a high concentration of lipids (Shahmoradian et al., 2019; Spillantini, Crowther, Jakes, Hasegawa, & Goedert, 1998). Five pathogenic point mutations in *SNCA*, which encodes alpha-synuclein, are known to cause autosomal dominant PD: p.A53T (Polymeropoulos et al., 1997), p.A30P (Krüger et al., 1998), p.E46K (Zarranz et al., 2004), p.G51D (Lesage et al., 2013), and p.A53E (Pasanen et al., 2014). Duplications or triplications of the *SNCA* locus also cause familial PD, indicating that increased levels of wildtype alpha-synuclein may be sufficient to cause disease (Chartier-Harlin et al., 2004; Singleton et al., 2003). Indeed, genome wide association studies (GWAS) show that single nucleotide polymorphism (SNP) genetic variants in *SNCA* are a risk factor in sporadic PD related to modulation of alpha-synuclein expression (Chiba-Falek, Lopez, & Nussbaum, 2006; Edwards et al., 2010; Pihlstrøm et al., 2018).

In addition to neuronal alpha-synuclein aggregates, numerous cases of alpha-synuclein positive astrocytes have also been identified in the post-mortem brain of idiopathic PD patients (Braak, Sastre, & Del Tredici, 2007; Song et al., 2009; Wakabayashi, Hayashi, Yoshimoto, Kudo, & Takahashi, 2000). Furthermore, glial and oligodendrocyte alpha-synuclein inclusions have been found in the post-mortem brain of patients with the p.A30P, p.A53T and p.G51D *SNCA* mutations (Kiely et al., 2013; Markopoulou et al., 2008; Seidel et al., 2010). One reason this pathology was only rarely described in earlier reports may be that astrocyte alpha-synuclein inclusions are poorly labelled or unlabeled when using N- and C-terminal antibodies, and can be visualized only using harsh antigen retrieval methods (Sorrentino, Giasson, & Chakrabarty, 2019). Taken together, these data indicate that glial pathology occurs in sporadic and familial PD, although alpha-synuclein glial aggregates/inclusions are likely to be underreported.

The motor symptoms of PD typically occur when approximately 70% of nigral neurons are degenerated (Fearnley & Lees, 1991). However, in the adult brain neurons exist in a dynamic microenvironment with glial cells at an approximate ratio of 1:1 (von Bartheld, Bahney, & Herculano-Houzel, 2016), with astrocytes being the most abundant cell type (Azevedo et al., 2009). Glia, and specifically astrocytes, regulate and support neurons at the tripartite synapse, are involved in synaptogenesis, synaptic pruning, neuronal branching, and the release and regulation of metabolites, neuro- and gliotransmitters and neurotrophic factors (Eroglu & Barres, 2010; Harada, Kamiya, & Tsuboi, 2016). This glial support is underpinned by transient elevations in astrocyte calcium (Ca^2+^) signalling, although this is still not fully understood (Bazargani & Attwell, 2016). Dysregulated calcium homeostasis has long been implicated in PD (Schapira, 2013), with increasing Ca^2+^ levels leading to clustering of alpha-synuclein at the synaptic vesicle (Lautenschläger et al., 2018).

The interplay between astrocytes and neurons in PD pathogenesis has become a focus of intensive research. Astrocytes exist in neurotoxic (A1) or neuroprotective (A2) subsets, and A1 astrocytes are found in abundance in the SNc of idiopathic PD patients (Liddelow et al., 2017). Astrocytes act as effective scavengers of alpha-synuclein, and the transmission of alpha-synuclein from neurons-to-astrocytes, astrocytes-to-astrocytes and astrocytes-to-neurons have all been reported (Cavaliere et al., 2017; H.-J. Lee et al., 2010; Rostami et al., 2017). In summary, there is emerging evidence pertaining astrocyte dysfunction in the pathogenesis of PD.

In this present study, we generated iPSC-derived midbrain astrocytes from patients carrying pathogenic mutations in *SNCA* and from healthy controls. Many human astrocyte culture protocols use commercial animal serum, which not only has seasonal and geographical batch-to-batch variation, but also contains a non-defined mixture of components that renders the astrocytes reactive (Gstraunthaler, Lindl, & Valk, 2013; Magistri et al., 2016; Perriot et al., 2018). We generated astrocytes using a novel serum-free protocol that supported astrocyte growth and maturity for a minimum of 140 days, expressing the midbrain marker FoxA2 in addition to astrocytic markers including ALDH1L1, Vimentin, AQP4, S100β and GFAP. In astrocytes derived from patients with *SNCA* mutations, we observed aggregation of alpha-synuclein and significant elevation in Ca^2+^ release. An in-depth assessment of mitochondrial morphology determined that mutant astrocytes have increased mitochondrial fragmentation contributing to mitochondrial dysfunction. Overall, our results directly indicate that astrocytes harbouring pathogenic *SNCA* mutations are dysfunctional, and intervention strategies for the rescue of non-neuronal cells should be considered in the early stages of the disease.

## Methods

### Ethics

Ethical approval for the development of and research pertaining to patient-derived cell lines was granted given by the National Ethics Board of Luxembourg, (Comité National d’Ethique dans la Recherche; CNER #201411/05).

### Cell lines

The induced pluripotent stem cell (iPSC) lines used in this study are shown in Table 1 with the culture conditions previously described (Simone B Larsen et al., 2020).

**Table 1:** Summary of patient-derived cell lines used in this study

| Cell name | Status | Gender | Age at Biopsy | Age of onset | Reprogramming method | Mutation |
| --- | --- | --- | --- | --- | --- | --- |
| Ctrl 16 | Control | Male | 72 | - | Sendai | - |
| Ctrl 17 | Control | Male | 67 | - | Lentivirus | - |
| Ctrl 18 | Control | Female | 72 | - | Sendai | - |
| A30P | PD | Male | 67 | 55 | Lentivirus | c.88G>C p.A30P *SNCA* |
| A53T | PD | Female | 51 | 39 | Sendai | c.157G>A p.A53T *SNCA* |
| Duplication | PD | Female | 67 | 48 | Sendai | *SNCA* gene locus duplication |
| Triplication | PD | Female | 55 | 50 | Sendai | *SNCA* gene locus triplication |

### Generation of neural precursor cells

The neural precursor cells (NPCs) were generated and cultured according to an established small molecule NPC (smNPC) protocol (Reinhardt et al., 2013). The characterisation of the homogenous NPCs is shown in Supplementary Figure 2 with the culture conditions detailed in the Supplementary Experimental Procedures.

### Generation of midbrain astrocytes

NPCs dissociated to single cells using Accutase (ThermoFisher) were plated 4×10^5^ in a well of a 6well plate pre-coated with Geltrex™ (ThermoFisher). The media used for the glial induction was a 50:50 mix of DMEM/F12 and Neurobasal, containing 1% B27, 0.5% N2, plus 1% Glutamax and Penicillin/Streptomycin (all Life Technologies). Cells were passaged in a ratio of 1:5 upon reaching confluence within the first 10 days of astrocyte patterning and proliferation; thereafter astrocytes were passaged at a ratio of 1:2. All experiments described in this manuscript were performed from day 60 onwards after the beginning of astrocyte differentiation.

### Generation of midbrain dopaminergic neurons

NPCs were differentiated to midbrain dopaminergic neurons according to the previously published protocol (Reinhardt et al., 2013).

### Real-time PCR

The RNA was extracted and reverse transcribed to cDNA as previously published (Simone B Larsen et al., 2020). RT-PCR was used to detect transcripts of the following genes (5’-3’): GFAP, forward: ATCCCAGGAGCGAGCAGAG, reverse: CCCAGCCAGGGAGAAATCCA; Aquaporin-4 (AQP4), forward: TGAGTGACAGACCCACAGCA, reverse: TTGATGGTGGATCCCAGGCTG; S100 calcium-binding protein β (S100β), forward: TGGAAGGGAGGGAGACAAGC, reverse: CCTGGAAGTCACATTCGCCG; Alpha-2-Macroglobulin (A2M), forward: AGCTTTGTCCACCTTGAGCC, reverse: CAGTTCGGACAATGCCTCCC; Tyrosine hydroxylase (TH), forward: AGTGCACCCAGTATACCGC, reverse: TCTCAGGCTCCTCAGACAGG; β-actin, forward: GAAGTTGGGTTTTCCAGCTAA, reverse: GGAGAACAATTCTGGGTTTGA. KOD Hot Start DNA polymerase (0.02U/μL; Merck) was used with the following programme: Pre-denaturation (95°C; 2mins), 35 cycles of denaturation (95°C; 30s), annealing (60°C; 45s) and extension (70°C; 60s), followed by a final extension (70°C; 5mins). The results were normalized to β-actin.

### Immunocytochemistry

Astrocytes were fixed at d120 of directed differentiation from NPCs with the fixation, permeabilization and imaging performed as previously detailed (Simone B Larsen et al., 2020). The primary antibodies used were anti-alpha-synuclein (1:250; BD Transduction, #610787), -Connexin-43 (1:150; Santa Cruz, sc-271837), -FoxA2 (1:100; Santa Cruz, sc-101060), -GI Syn (1:100; Santa Cruz, sc-74430), -GFAP (1:1000; Dako, Z0334), -NF68 (1:200; Sigma, N-5139), -S100β (1:100; Abcam, ab868) –TUBB3 (1:300; Covance, MRB-435P) -Vimentin (1:100; Santa Cruz, sc-373717). Secondary antibodies used were: Alexa Fluor 488 Goat anti-Mouse IgG (H+L) (1:1000; Invitrogen, A11029), Alexa Fluor 568 Goat anti-Rabbit IgG (H+L) (1:1000; Invitrogen, A11036). Hoechst 33342 (Invitrogen, H3570) was used as a nuclei counterstain.

### Cell counts

NPCs were directly differentiated to midbrain astrocytes for over 80 days, plated onto coverslips and fixed and stained for the neuronal marker TUBB3, the pan-astrocytic marker Vimentin, and the nuclear stain Hoechst 33342. Between 5-10 fields of view (FOV) were acquired per coverslip using the 10-25x objective lens with approximately 1500 total cells counted. The merged image was processed in ImageJ where the image was converted to RGB stack, with the three stains shown in separate (red/green/blue) channels. The “Cell Counter” plug-in was used to count the Hoechst-stained nuclei before the TUBB3 stained neurons were quantified on the overlaid image.

### Protein Immunoblotting

The protein immunoblotting cells pellets were extracted using RIPA buffer (Tris HCl pH 7.4, (50 mM); NaCl (150 mM); Triton-X-100 (1%); sodium deoxylcholate (0.5%); SDS (0.1%); EDTA (1mM); Tris-HCl (50mM); NaCl (150mM)) plus 1 tablet of cOmplete™ proteinase inhibitor cocktail (Roche) per 20mL of RIPA buffer. Polyacrylamide gels (10%) were blotted by dry transfer on 0.2μM nitrocellulose membranes using the iBlot™ 2 Gel Transfer Device (ThermoFisher). Primary anti-alpha-synuclein (1:1000; BD Transduction, #610787) and -β-actin (1:20,000; Cell Signaling, 3700S) antibodies were probed overnight followed by incubation for 1 hour with secondary antibodies conjugated to HRP (Invitrogen). Densitometry was performed using ImageJ (Schneider, Rasband, & Eliceiri, 2012) and normalized to β-actin.

### Flow Cytometry

Astrocytes were dissociated to single cells using Accutase and fixed dropwise whilst vortexing in 4% PFA before being placed onto an orbital roller for 15 minutes. Cells were incubated in Saponin buffer (0.05% Saponin, 1% BSA). The primary antibodies used were anti-ALDH1L1 (1:100; Santa Cruz, sc-100497), GFAP (1:300; Dako, Z0334), S100β (1:100; Abcam, ab868). The secondary antibodies used are detailed above. Secondary antibody only-stained cells were used as a gating control. Flow cytometry was run using the LSRFortessa™ (BD Biosciences) cell analyzer with FlowJo software version 10.0.7 (LLC) used for the visualizations of the graphs.

### Detection of Calcium

5×10^4^ astrocytes were seeded onto 8 well chamber slides coated with polyornithine and laminin and left for 48 hours. The Fluo-4 Calcium Imaging Kit (Invitrogen™, F10489) was used for the detection of cytosolic Ca^2+^ according to the manufacturer’s guidelines. For the live cell image acquisition, live-streaming mode was used for 20 minutes per cell line on a pre-heated stage set to 37°C within a heated chamber. The Zen 2.3 software (blue edition) on a Zeiss Spinning Disk confocal microscope (Carl Zeiss Microimaging GmBH) was used for the acquisition. For the detection of spontaneously released cytosolic calcium, the video was exported as single image files and a minimum of 50 firing cells per cell line were selected using Fiji (Schindelin et al., 2012). The calcium signaling analyzer (CaSiAn) tool was used to analyze the Ca^2+^ spikes and is previously described (Moein et al., 2018).

### Measurement of oxygen consumption rate

Oxygen consumption rate (OCR) and extracellular acidification rate (ECAR) were measured from the plated astrocytes using the Seahorse XFe96 Cell Metabolism Analyzer (Agilent). Astrocytes were dissociated to single cells and plated at a density of 1×10^4^ per well, a minimum of 6 wells were used per line per experiment. Mitochondrial respiration was determined using a mitochondrial stress test 48 hours after plating according to the manufacturer’s instructions using the Seahorse Wave software (2.6.0, Agilent). The final concentrations of Oligomycin (1μM; #75351), FCCP (500nM; #C2920), Rotenone (5μM; #R8875) and Antimycin A (5μM; #A8674) (all Sigma) used in the stress test were normalized by total protein. This was performed directly after the conclusion of the stress test by lysing with RIPA buffer and calculating the total protein using a BCA assay.

### Assessment mitochondrial morphology and pyknotic nuclei

Mitochondrial morphology was assessed by Tom20, a subunit of the translocase of the mitochondrial outer membrane (TOM) complex. At day 60 of astrocyte differentiation, 5×10^4^ astrocytes were seeded onto 8 well chamber slides coated with polyornithine and laminin and left for 48 hours. The astrocytes were fixed in paraformaldehyde (4% in PFA) and permeabilized using Triton-X-100 as previously described (Simone B Larsen et al., 2020). The mouse anti-Tom20 antibody (1:1000; Santa Cruz, sc-17764) was used with the Alexa Fluor 488 (Life Technologies) antibody used to detect Tom20. Hoechst 33342 (Invitrogen; #H3570) was used as a nuclear stain and for normalization. A minimum of ten aleatory fields were acquired for each condition using the 63x objective as Z-stacks (at 0.26μm intervals) covering the entire depth of the cell. Images were acquired using the Zeiss Spinning Disk confocal microscope (Carl Zeiss Microimaging GmBH). Maximum intensity projection was used to visualize the 3D Z-stacks for the 2D analysis of mitochondrial morphology. The morphometric analysis of the mitochondrial morphology as shown in Table 2 was previously established with morphometric features defined (Antony et al., 2020). Briefly, Form Factor = Perimeter^2^/4πArea; Aspect Ratio = Major axis length/minor axis length; Mitochondrial number = Mitochondrial pixels / nuclei pixels. MitoPerimeterProportion_Norm = MitoPerimeterPixels / MitoPixels; MitoShapeByPerimeter_Norm = MitoBodyPixels/MitoPerimeterPixels; MitoSkelProportion_Norm = MitoSkel/MitoPixels; MitoErosionBodies_Norm = MitoBodies/MitoCount; PyknosisMetric_Norm = PyknoticPixels/NucleiPixels.

**Table 2:** Summary of mitochondrial morphology and cell death analysis in the patient-derived astrocytes harbouring mutations in *SNCA*. A minimum of ten aleatory fields per replicate were acquired for each condition using the 63x objective as Z-stacks. The obtained images were analyzed using a Matlab script capable of analyzing 3D nuclear staining and 3D mitochondrial networks that has been previously reported (Antony et al., 2020). Statistical analysis was performed using a one-way ANOVA with Dunnett’s multiple comparison post-hoc test. ****p<0.0001, ***p<0.0005, **p<0.001 *p<0.05.

|  | Patient-derived Astrocytes DIV90 | Direction vs controls | A30P | A53T | Duplication | Triplication |
| --- | --- | --- | --- | --- | --- | --- |
| Fragmentation 2D | Mean_FormFactor | Decreased | *** | **** | ** | ** |
|  | Mean_Aspect Ratio | Decreased |  | **** | **** | **** |
|  | Mitochondrial number | Decreased |  | **** |  | ** |
| Fragmentation 3D | MitoPerimeterProprtion_Norm | Decreased | *** | ** | * | *** |
|  | MitoShapeByPerimeter_Norm | Increased | *** | ** | * | *** |
|  | MitoSkelProportion_Norm | Decreased |  | *** | * | * |
|  | MitoErosionBodies_Norm | Decreased |  | * | * | **** |
| Pyknosis | PyknosisMetric_Norm | Increased |  | **** | * | *** |

### Statistical analysis

GraphPad Prism^®^ (Version 8.3.0, GraphPad Software Inc., USA) was used for statistical analyses. The type of statistical analyses performed and P-value for each experiment can be found in the legend of each figure.

## Results

### Generation of patient-derived midbrain astrocytes using a serum-free protocol

We used a previously published protocol to generate rapidly expandable homogenous neural precursor cells (NPCs) from patient-derived iPS cells lines (Table 1) (Reinhardt et al., 2013). The NPCs were cryopreservable and committed as a multipotent progenitor to the neural lineage without pluripotent or non-neural contamination, and were used as a starting point for astrocyte differentiation (Figure 1A). Astrocyte-specific cultures were successfully generated from three healthy control iPSC lines and four *SNCA* mutant iPSC lines: the point mutations p.A30P and p.A53T, and the duplication and triplication of the *SNCA* gene locus. For regionally-specific astrocyte specification and patterning, the midbrain-hindbrain boundary transcription factor fibroblast growth factor (FGF) 8b (100ng/mL; Peprotech, 100-25) was used to promote midbrain identity (S. M. Lee, Danielian, Fritzsch, & McMahon, 1997). Epidermal growth factor (EGF)(20ng/mL; Peprotech, AF-100-15) and FGF2 (10ng/mL; Peprotech, 100-18B) was used to specify gliogenesis (Liu & Neufeld, 2007), with Heparin (5µg/mL; Sigma, H3149) added to potentiate the effect of FGF2 (Caldwell, Garcion, terBorg, He, & Svendsen, 2004). Leukaemia inhibitory factor (LIF)(5ng/mL; Peprotech, AF-300) was added to activate the JAK-STAT pathway committing the cells to gliogenesis (Bonni et al., 1997), and the histone deacetylase (HDAC) inhibitor Valproic acid (VPA)(1mM; Sigma, P4543) was used to increase the expression of GDNF (Rincón Castro, Gallant, & Niles, 2005) and glial precursor proliferation (H. J. Lee, Dreyfus, & DiCicco-Bloom, 2016). The mitogens FGF2, FGF8b and EGF along with Heparin, LIF and VPA were removed from the culture to induce the immature proliferative astrocytes to differentiate into terminally differentiated astrocytes. Heregulin 1β (5ng/mL; Peprotech, 100-03) and CNTF (5ng/mL; Peprotech, 450-13) was used to induce the differentiation of the precursors, with Heregulin 1β removed at day 30 of directed astrocyte differentiation (Pinkas-Kramarski et al., 1994). The committed astrocytes were left to mature in terminal differentiation media with CNTF maintaining the upregulated JAK-STAT pathway (Bonni et al., 1997). After 30 days of directed astrocyte differentiation the immature astrocytes from the Ctrl 17 line were >80% GFAP positive, 30-70% S100β positive, and <5% positive for ALDH1L1. Continued maturation for 90 days increased the proportion of both GFAP and S100β positive cells to >98%, and the percentage of ALDH1L1 positive cells increased with time to 50-70% (Figure 1B).

**Figure 1:**
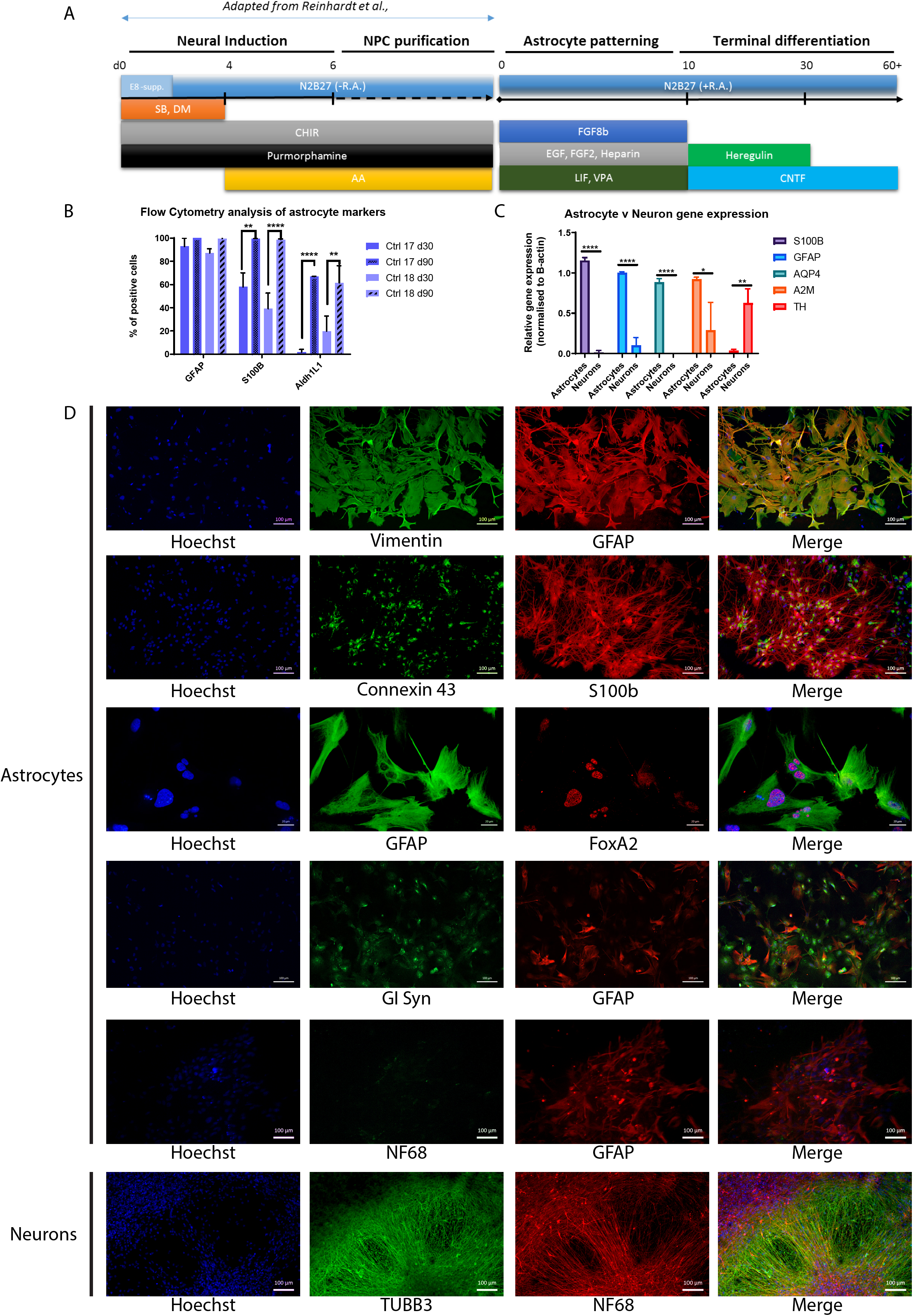
Differentiation and characterisation of midbrain astrocytes from human iPSCs via a NPC state. **(A)** An overview of the NPC generation and directed astrocyte differentiation protocol. **(B)** Quantitative analysis of astrocyte marker proteins using flow cytometry. Astrocytes were differentiated for 30 and 90 days. A 2-way ANOVA was used with Tukey multiple comparison post-hoc test: ****p<0.0001, ***p<0.0002. **(C)** Relative gene expression of astrocytes normalized to B-actin. Astrocytes were differentiated for 120 DIV; the neurons were differentiated for 45 DIV according to Reinhardt et al. (Astrocytes, n=3 cell lines; Neurons, n=4 cell lines; an unpaired t test was used with a two-tailed P value: ****p<0.0001, **p=0.0025, *p=0.0272 **(D)** Triple immunofluorescence labelling of astrocytes expressing pan astrocytic and midbrain specific markers. Astrocytes were differentiated for 140 days with representative images shown.

At d120, we compared astrocyte gene expression to d45 iPS-derived midbrain dopaminergic neurons generated using a previously published protocol (Reinhardt et al., 2013)(Figure 1C). The expression of the astrocyte-specific genes *S100β, GFAP, AQP4, A2M* were all significantly higher in astrocytes than neurons, and the rate-limiting enzyme in catecholamine biosynthesis that converts tyrosine to L-Dopa and specifically found in dopaminergic neurons was enriched in neurons compared to the astrocytes (Figure 1C). Immunocytochemistry revealed robust expression of GFAP, S100β, Vimentin, Connexin 43, Glutamine Synthetase (GI Syn), the regionally specific midbrain marker FoxA2 (Figure 1D) and the absence of neurofilament 68 protein (NF68). Astrocyte purity was quantified by determining the number of cells expressing the pan-astrocytic marker Vimentin and the number expressing the neuronal marker β-3-Tubulin (TUBB3). Each astrocyte line was >90% pure, and there was no significant difference in purity between controls and astrocytes harbouring *SNCA* mutations (Supplementary Figure 1).

### Aggregation of alpha-synuclein in astrocytes harbouring *SNCA* mutations

Alpha-synuclein protein was detectable in all astrocyte lines, and highest in the *SNCA* triplication cell line, although approximately 10-fold lower than the level detectable in iPS-derived neurons (Figure 2A). A low but detectable population of contaminating neurons were found in these cultures (Supplementary Figure 1). Immunocytochemistry confirmed the presence of alpha-synuclein in GFAP^+^ astrocytes (Figure 2B). In healthy control lines, alpha-synuclein was found well distributed and typically localised to the cytoplasm, whereas alpha-synuclein aggregates were detectable in astrocytes harbouring the pathogenic *SNCA* mutations (Figure 2B).

**Figure 2:**
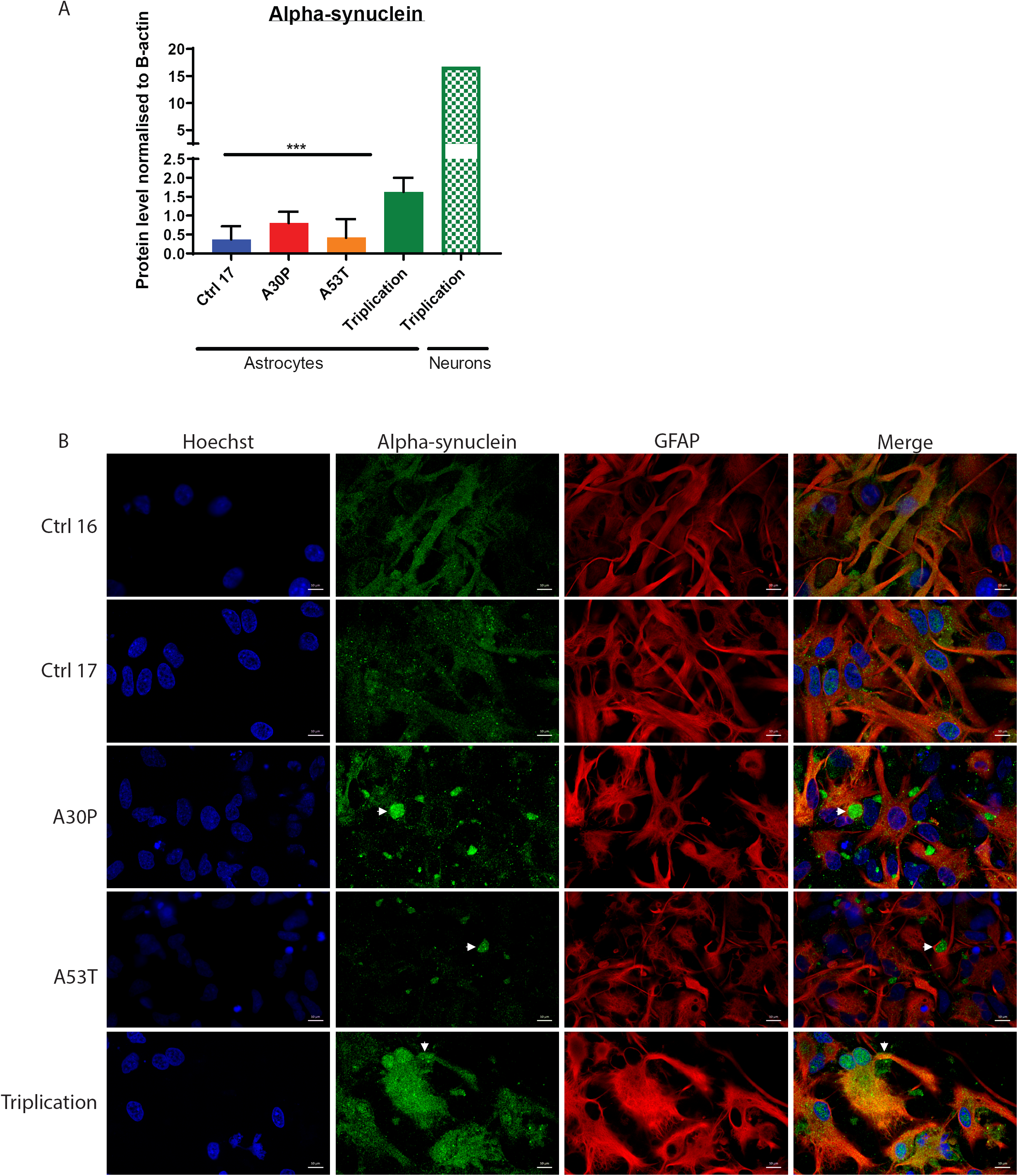
Altered distribution of alpha-synuclein in patient-derived astrocytes. **(A)** Normalized immunoblot data show detectable level of alpha-synuclein in patient-derived astrocytes. **(B)** Immunostaining showing colocalisation of alpha-synuclein protein with GFAP^+^ astrocytes in the control and patient cell lines. The white arrowheads in the A30P, A53T and Triplication astrocytes samples show aggregated alpha-synuclein.

### Excess cytosolic Ca^2+^ release in *SNCA* mutant astrocytes

To validate that the astrocytes were functionally mature, we assessed the physiological propagation of intercellular Ca^2+^ waves using the Fluo-4 AM indicator of cytosolic Ca^2+^ under basal conditions. We were able to detect calcium waves in all cell lines and used the Calcium Signal Analyzer (CaSiAn) software tool (Moein et al., 2018) to quantify the Ca^2+^ dynamics (Figure 3A-3B). Astrocytes harboring p.A30P and p.A53T point mutations or the *SNCA* triplication had increased amplitude of Ca^2+^ spikes (Figure 3C), indicating increased levels of cytosolic Ca^2+^ compared to the control lines. Additionally, the p.A30P and p.A53T lines released Ca^2+^ into the cytoplasm at greater rates than the controls (Figure 3D), and astrocytes with the *SNCA* triplication line had a greater spike triangle (Figure 3E), indicating increased Ca^2+^ release per Ca^2+^ spike.

**Figure 3:**
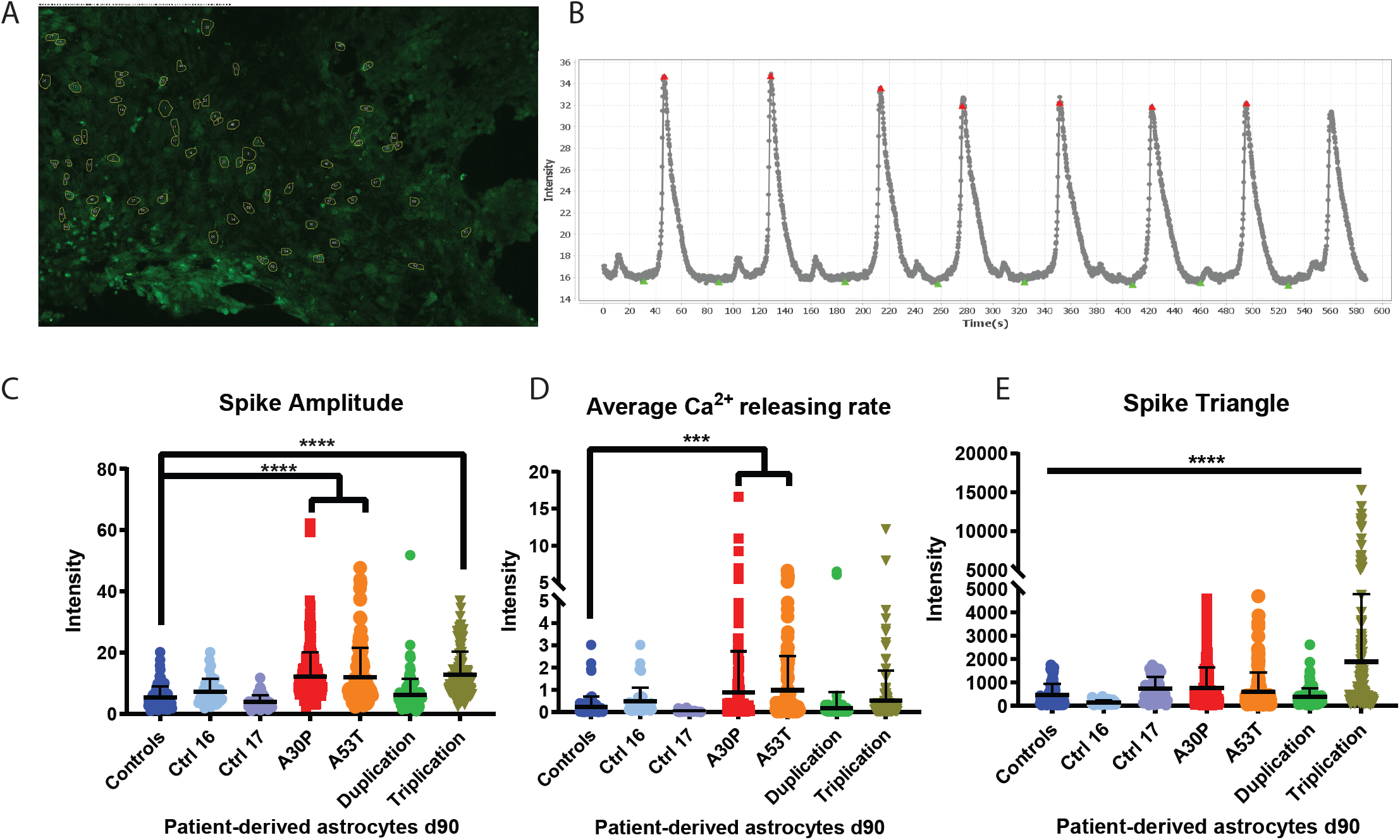
Assessment of cytosolic Ca^2+^ in patient-derived midbrain astrocytes harbouring *SNCA* mutations. **(A)** Calcium waves recorded over 10 minutes at 37°C using the cytosolic calcium tracer Fluo-4 AM at d90 of differentiation (n=3). **(B)** The Calcium Signal Analyser software (CaSiAn) was used to quantify the calcium waves with the red triangles corresponding to the identified peaks and the green triangles the nadirs. The measurable parameters: **(C)** Spike amplitude, **(D)** Average calcium releasing rate, and **(E)** Spike triangle were quantified using an unpaired one-way ANOVA with Dunnett’s multiple comparison post-hoc test: ****p<0.0001, ***p<0.0005. All data expressed as mean ± SD.

### Astrocytes containing pathogenic mutations in *SNCA* have fragmented mitochondria and increased cell death

In many cellular models of PD, excess or pathogenic alpha-synuclein has been shown to lead to mitochondrial fragmentation in neurons (Guardia-Laguarta et al., 2014; Kamp et al., 2010; Nakamura et al., 2011; Zambon et al., 2019). Additionally, increased abundance of alpha-synuclein leads to increased transfer of Ca^2+^ from the ER to the mitochondria (Calì, Ottolini, Negro, & Brini, 2012), and Ca^2+^ overload has been associated with mitochondrial fragmentation and cell death (Granatiero, Pacifici, Raffaello, De Stefani, & Rizzuto, 2019). Therefore, we assessed mitochondrial form factor and aspect ratio in the patient-derived astrocytes. At d90 of differentiation, lines harbouring *SNCA* mutations showed increased mitochondrial fragmentation compared to control astrocytes (Figure 4A-C), similar to findings in PD neurons (S B Larsen, Hanss, & Krüger, 2018). Additional in-depth mitochondrial morphometric analysis of 3D mitochondrial networks (Antony et al., 2020) revealed an increased number of swollen, spherical mitochondria with reduced branching and connectivity in the *SNCA* mutant astrocytes compared to controls (Table 2). Notably, the lines harbouring the two *SNCA* mutations that cause earlier disease onset and increased clinical severity—p.A53T and *SNCA* triplication—showed more severe mitochondrial morphological impairment than *SNCA* duplication line, indicating an alpha-synuclein dosage effect. The p.A30P mutation line had deficits in mitochondrial branching (form factor), whereas mitochondrial length (aspect ratio) and number was unaffected. Lastly, we determined the presence of pyknotic nuclei, an early indicator of cell death. The p.A53T and *SNCA* triplication lines had an increased number of pyknotic nuclei, there was a mild elevation in the number of pyknotic nuclei in the *SNCA* duplication line, and no difference between the p.A30P and controls (Table 2).

**Figure 4:**
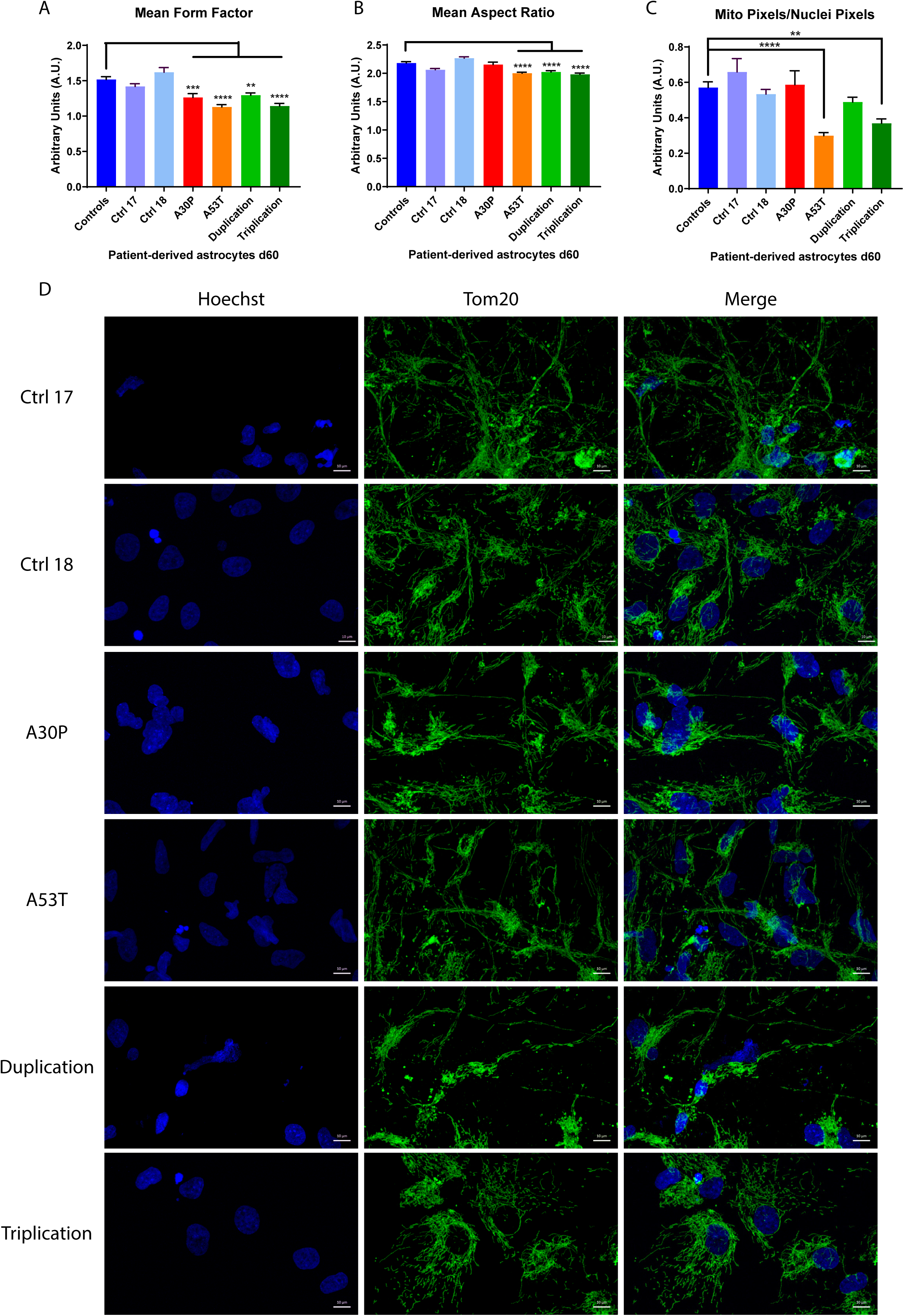
Assessment of fragmented mitochondria in astrocytes harbouring pathogenic mutations in SNCA after 90 days of directed differentiation (n=3). Patient-derived astrocytes containing mutations in *SNCA* have more fragmented mitochondria than the unaffected controls assessed by **(A)** Form factor, **(B)** Aspect ratio and **(C)** Mitochondrial number. A minimum of ten aleatory fields were acquired for each condition using the 63x objective as Z-stacks (n=3). The obtained images were analysed using an in-house developed Matlab script capable of analysing mitochondrial 3D networks and 3D nuclear staining. **(D)** For assessment of 2D fragmentation parameters, a maximum intensity projection was used with a representative example for each cell line shown. Statistics was performed using a one-way ANOVA with Dunnett’s multiple comparison post-hoc test: **p<0.001, ****p<0.0001.

### Aberrant mitochondrial function in heterozygous mutant *SNCA* astrocytes

We evaluated the bioenergetic profile of the patient-derived astrocytes by assessing mitochondrial function using oxygen consumption rate (OCR) and extracellular acidification rate (ECAR) during a mitochondrial stress test. Astrocytes harbouring p.A30P and p.A53T heterozygous mutations had significantly reduced OCR compared to the controls (Figure 5A) and the p.A53T and triplication mutants had significantly reduced ECAR compared to controls (Figure 5B). Furthermore, both the p.A53T line and the pA30P line exhibited deficits in maximum respiration and non-mitochondrial oxygen consumption, and astrocytes harboring the p.A53T alpha-synuclein mutation had deficient spare respiratory capacity (Figures 5C-H).

**Figure 5:**
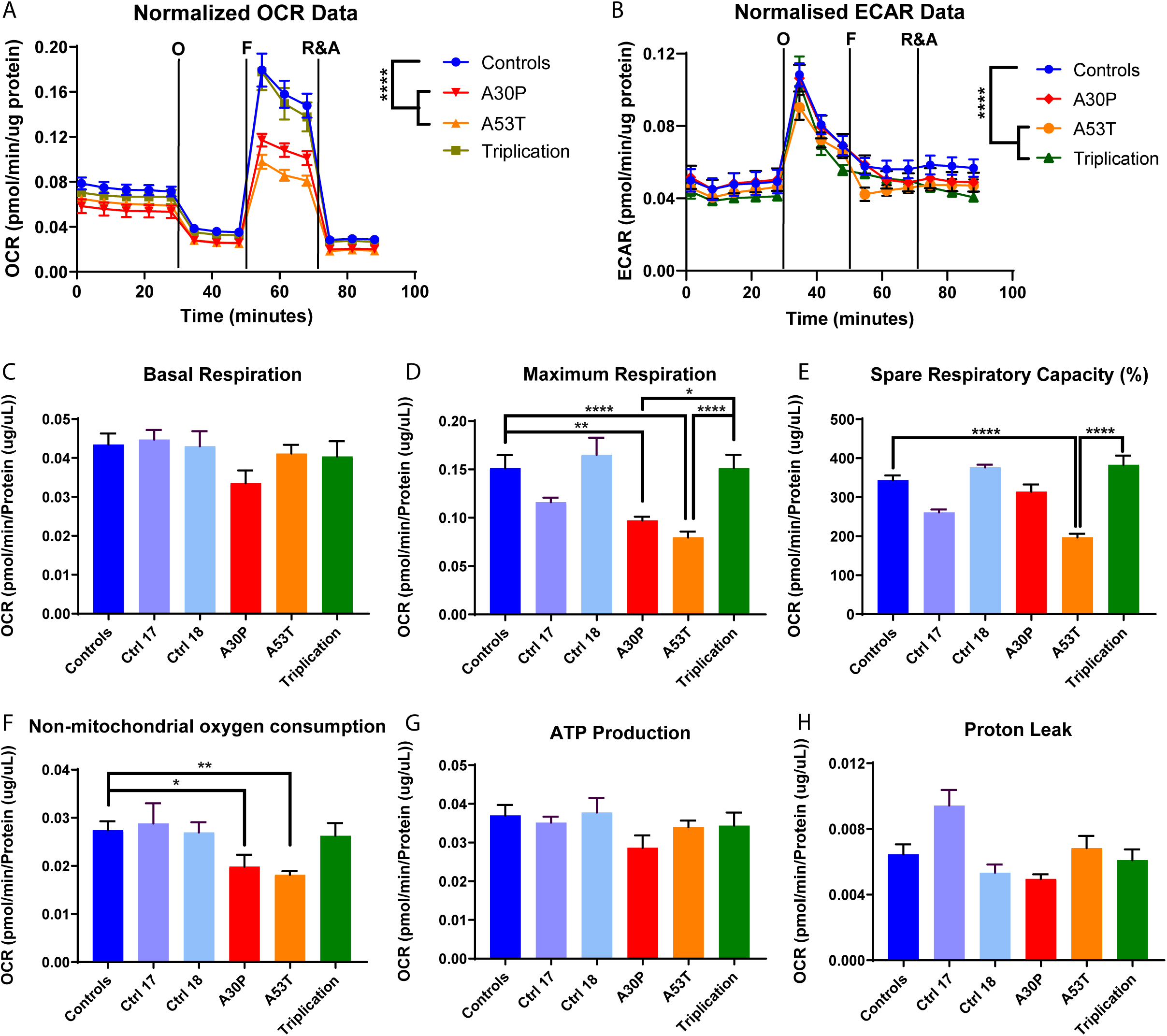
Mitochondrial bioenergetics of patient-derived astrocytes harbouring pathogenic mutations in *SNCA* undergoing a mitochondrial stress test. Detection of mitochondrial respiration via **(A)** oxygen consumption rate (OCR) or **(B)** extracellular acidification rate (ECAR) under basal conditions and following the treatments of the ATP synthase inhibitor Oligomycin (O, 1uM), the oxidative phosphorylation uncoupler FCCP (F, 500nM), and the electron transport chain inhibitors Rotenone (Complex I) and Antimycin A (Complex III) (R&A, 10uM). The cumulative OCR or ECAR profile is shown of astrocytes differentiated for 60 days of differentiation (n=3). The combined profiles are plotted as mean for visual clarity. A two-way ANOVA was performed with Dunnett’s post-hoc test: ****p<0.0001,*p=0.0124. The rates of **(C)** basal respiration, **(D)** maximum respiration **(E)** spare respiratory capacity (%), **(F)** non-mitochondrial oxygen consumption, **(G)** ATP production and **(H)** proton leak were calculated using a one-way ANOVA using Tukey post-hoc test. ****p<0.0001, ***p<0.002, **p<0.005, *p<0.001.

## Discussion

Astrocytes are heterogeneous and display inter- and intra-regional distinctions (Morel et al., 2017). Previous studies have generated region-specific astrocytes from human iPS cells representing the spinal cord (Roybon et al., 2013) and the forebrain (Bradley et al., 2019). We developed a serum-free protocol that for the first time shows the generation of midbrain astrocytes from patient-derived iPSCs, which have particular relevance for the study of PD. This protocol notably results in enrichment in the astrocyte maturation marker ALDH1L1 over time, and expression of the midbrain specific marker FoxA2, the pan-astrocytic marker Vimentin, mature astrocyte markers Connexin 43, GI Syn and *AQP4*, and the commonly used astrocyte markers S100β and GFAP.

The level of *SNCA* expression in astrocytes compared to neurons and oligodendrocytes is low (Booth, Hirst, & Wade-Martins, 2017; Zhang et al., 2016). Although there is considerable evidence of alpha-synuclein accumulation in astrocytes (Braidy et al., 2013; Cavaliere et al., 2017; di Domenico et al., 2019; Rostami et al., 2017), it is generally accepted that astrocytes themselves do not express detectable levels of the soluble monomeric alpha-synuclein protein. We speculate that in our system, the presence of a small but detectable population of neurons generates alpha-synuclein that is subsequently taken up by the astrocytes.

During aging and disease, astrocytes display Ca^2+^ dysregulation, characterized by extensive cytosolic Ca^2+^ levels, increased Ca^2+^ transients and more frequent Ca^2+^ oscillations (Kuchibhotla, Lattarulo, Hyman, & Bacskai, 2009). In our model of patient-derived iPS-derived midbrain astrocytes, we show dysregulation of cytosolic Ca^2+^ in the astrocytes carrying pathogenic *SNCA* mutations. However, the mode of astrocyte dysregulation in the form of Ca^2+^ overload differs between the *SNCA* triplication line and the heterozygous mutants. The astrocytes carrying the triplication of the *SNCA* gene released more cytosolic Ca^2+^ over a longer duration whereas both the A30P and A53T point mutations release excess Ca^2+^ due to increased rate of Ca^2+^ influx into the cytosol. Interestingly, the *SNCA* duplication line displayed no measurable differences in cytosolic Ca^2+^ compared to the controls, suggesting a critical threshold of physiological alpha-synuclein is required for Ca^2+^ cytosolic overload.

To date, this study is the most comprehensive assessment of PD patients harbouring different pathogenic *SNCA* mutations expressing physiological levels of the endogenous alpha-synuclein protein. Moreover, our analysis allows cross-correlations between the pathogenic mutations that have not been described before. Our data show that mitochondrial fragmentation is not only a key indicator of disease pathology found across different cellular models (Antony et al., 2020; S B Larsen et al., 2018; Mortiboys et al., 2008), but may also be an indicator of disease severity. The astrocytes carrying the A53T mutation have the most severe phenotypes across all of the parameters analyzed in this study. In accordance with this finding, individuals with the A53T mutation manifest PD symptoms approximately a 10-years earlier than carriers of other missense mutations (Kasten & Klein, 2013). Similarly, carriers of the *SNCA* triplication exhibit onset of PD symptoms earlier than carriers of *SNCA* duplication, and have more rapid disease progression (Kasten & Klein, 2013). Thus, the increased mitochondrial fragmentation, pyknotic nuclei and excess of cytosolic calcium found may all be biomarkers of *SNCA* dosage. Interestingly, the p.A30P line had reduced mitochondrial branching (form factor) yet no difference in mitochondrial length (aspect ratio), similar to findings found in idiopathic and familial PD fibroblasts (Antony et al., 2020; Mortiboys et al., 2008); which may indicate that loss of mitochondrial branching precedes loss of mitochondrial length and can serve as an indicator of disease severity.

The effects of PD mutations on neuronal phenotypes have been studied in patient-derived neurons, but it remained unclear if these mutations exerted pathological effects on other cell types. Our data show that astrocytes also possess measurable pathogenic phenotypes in response to PD-causing mutations that can contribute to the disease pathogenesis. This raises the hypothesis that in the brain, where neurons and non-neuronal support cells exists in a ratio of 1:1 (Azevedo et al., 2009), dysfunctional glia facilitate neuronal dysfunction. Furthermore, treatment of dysfunctional glia should be considered as an intervention strategy in the early stages of PD.

## Acknowledgments

The authors would like to thank the Parkinson’s disease patients and healthy control individuals that supported our research. Moreover, we thank Grazia Iannello and Susanna Usmmaa for their work on previous iterations of the astrocyte protocol. We thank Kathyrn Claiborn for reviewing the manuscript. We would like to acknowledge that Servier Medical Art (www.servier.com) created the figures in the graphical abstract, licensed under a Creative Commons Attribution 3.0 Unported License. The research was funded by grants from the Fonds National de la Recherche within the PEARL programme (FNR/P13/6682797), the National Centre for Excellence in Research on Parkinson’s disease (NCER-PD) programme, by the European Union’s Horizon 2020 research and innovation programme under Grant Agreement No 692320 (WIDESPREAD; CENTRE-PD) and by the National Institutes of Health (RF1 AG058475).

## Author contributions

Conceptualization: P.B. (Peter Barbuti) and R.K. (Rejko Krüger); methodology: P.B.; software: P.A. (Paul Antony); validation: P.B., F.M. (Francois Massart); formal analysis: P.B.; investigation: P.B.; resources: P.B., R.K., Gabriella Novak (G.N.), Simone Larsen (S.L.), Clara Berenguer-Escuder (C.B.), Bruno Santos (B.S.), Dajana Grossmann (D.G.), Takahiro Shiga (T.S.), Kei-ichi Ishikawa (K.I.), Wado Akamatsu (W.A.), Nobutaka Hattori (N.H.), and Steven Finkbeiner (S.F.); data curation: P.B.; writing—original draft preparation: P.B.; writing—review and editing: P.B., S.F. and R.K.; visualization: P.B.; supervision: R.K.; project administration: P.B. and R.K.; funding acquisition: R.K. All authors have read and agreed to the published version of the manuscript.

## Data availability statement

The data that support the findings of this study are available on request from the corresponding author. The data are not publicly available due to privacy or ethical restrictions.

