## Supplementary figures and images for "IPSC-derived midbrain astrocytes from Parkinson’s disease patients carrying pathogenic *SNCA* mutations exhibit alpha-synuclein aggregation, mitochondrial fragmentation and excess calcium release"

### Supplementary Figure 1

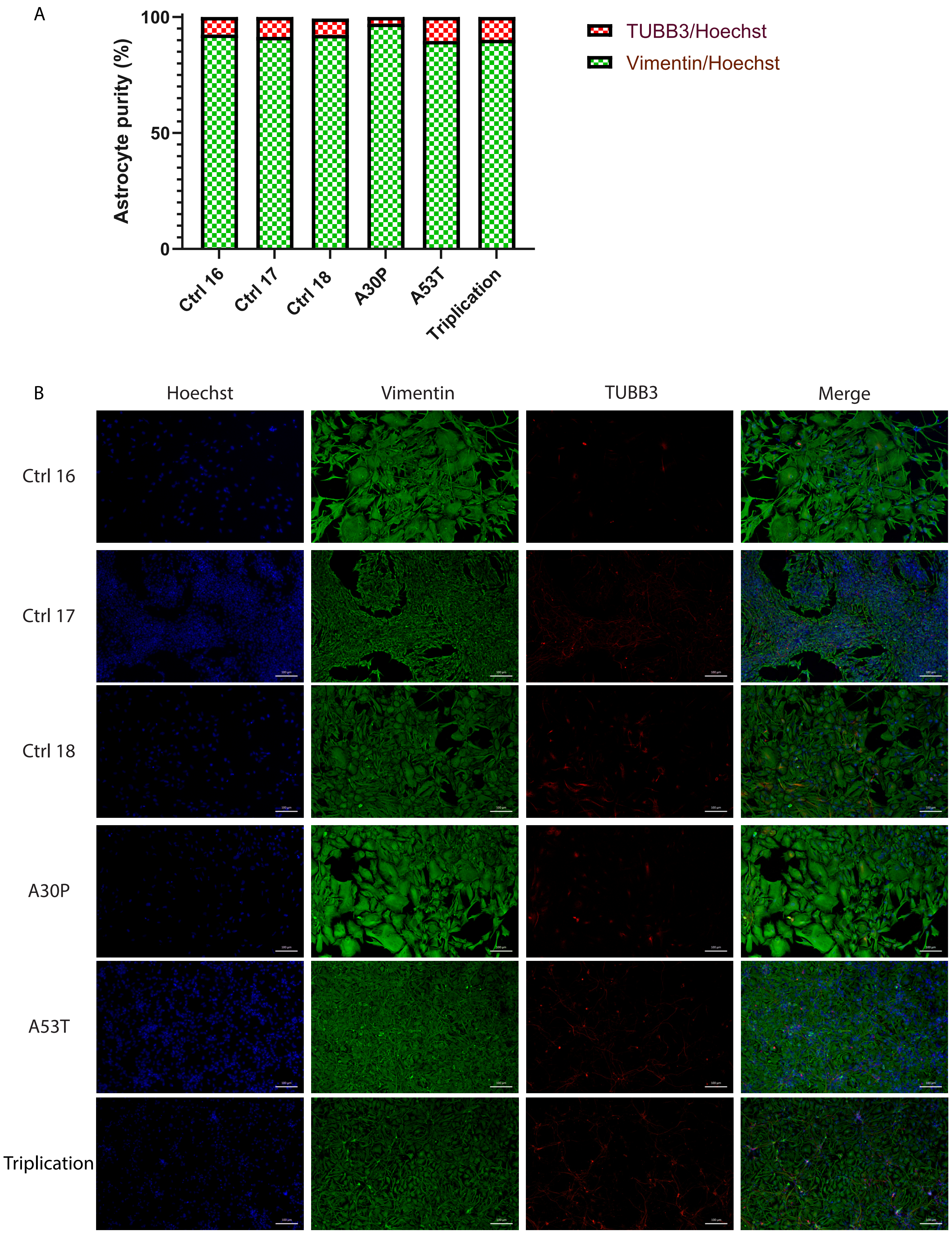
